# Modelling cancer cell dormancy in mice: statistical distributions predict reactivation times and survival dynamics

**DOI:** 10.1101/2025.03.21.644537

**Authors:** Nikolaos Sfakianakis, Dimitrios Katsaounis, Mark A.J. Chaplain

## Abstract

In predicting metastatic potential and improving treatment outcomes in cancer research, it is crucial that we understand the dynamics of cancer cell dormancy and reactivation. In this paper we propose, study, and evaluate a cancer growth model that incorporates cell death, dormancy, reactivation, and proliferation in the secondary sites. Using experimental data on mice, we test various statistical distributions and identify models that represent the asymmetry and variability observed in dormancy durations. Notably, the estimated cancer cell death rate remained consistent across all tested distributions, supporting its biological relevance as a robust parameter for modelling dormancy survival dynamics. The most suitable among the distributions we studied, the most suitable ones exhibit heavy-tails and asymmetric skewness; this aligns with the prolonged and rare dormancy periods expected of cancer cells. Our findings stress the importance of selecting appropriate statistical models for dormancy, both in predicting cancer cell reactivation events, and in informing therapeutic strategies that focus dormancy-driven metastasis.

## 1. Introduction

Cancer cell dormancy is an important stage in cancer progression, during which cancer cells pause/halt to proliferate but retain the possibility to reactivate in the future, a factor that plays a vital role in the spread of metastases [1, 9, 20]. The process is controlled by a complex network of biological processes; these include the tumour microenvironment, signals that are received and emitted by cells, various control mechanisms, and genetic and epigenetic changes [1, 15, 19]. Together, these dictate the period of dormancy of cancer cells and the conditions under which they can re-emerge to resume the disease process [2, 9].

As dormancy is stochastic in nature, it is essential to accurately measure the length of such periods in different populations of cancer cells. This is vital to assess the risk of metastasis and guide effective therapy [1, 2]. Intrinsic cellular mechanisms and extrinsic conditions within the tumour microenvironment, including nutrient availability and the proximity to blood vessels, influence such periods of dormancy [9, 16]. Thus, these intervals are very heterogeneous, and their trend is best characterized by the application of probability distributions echoing the biological heterogeneity seen in cell populations. It is essential to identify these trends to ascertain how frequently dormant cells re-activate and at what survival levels.

Mathematical modelling plays a pivotal role in analysing these dynamics [1, 16]. It allows scientists to compare different statistical distributions to empirical experimental data to determine the models that most accurately describe the behaviour of cancer cells under dormancy and reactivation. By comparing distributions that are well known to be asymmetric with long tails and duration variability, scientists are in a position to determine if dormancy tends to follow a simple exponential trend or more complicated models like Weibull, Gamma, Pareto, or Inverse Gaussian distributions. These tests help reveal variation in the probability of cells reactivating across time [6, 9].

This paper investigates a cancer growth model that addresses cancer cell death, dormancy, reactivation, and proliferation focusing on identifying the statistical distributions that best capture dormancy duration and reactivation times. We examine a variety of statistical distributions with a view to determining those which not only match the statistical data but are also consistent with biological facts [4]. Our findings form a basis for predictive models of cancer dormancy, our understanding of metastatic risks, and the development of new treatment protocols.

## 2 Insights into Cancer Cell Dormancy

The dormancy of cancer cells, particularly in the lung, is a highly critical phase of tumour growth and metastasis. Dormant cells are capable of remaining in a viable but non-proliferative state for extended periods of time, evading therapeutic treatment and ultimately leading to cancer recurrence. A number of factors, cellular, environmental, intra- and extratumoural are responsible for the regulation of the dormancy within the community of cancer cells. Here, we mention but a few:

### Extracellular Matrix (ECM) Interactions

Dormant cancer cells often exhibit strong adhesion to the ECM, which transmits biochemical signals that sustain their quiescent state. The ECM serves as a protective niche, maintaining dormancy by inhibiting cell proliferation and reinforcing cellular adhesion mechanisms. This interaction is critical in stabilizing the dormant phenotype and preventing reactivation [2, 9].

### Oxygen and Nutrient Availability

Avascular, in particular, tumour microenvironments can be characterized by hypoxia and limited nutrient availability. These environments possess the potential of cancer cell dormancy induction. Hypoxia is, for example, able to induce reduced metabolic activity and proliferation, while nutrient deprivation forces cells to resort to energy conservation status. These environmental stress conditions contribute to the likelihood of reactivation upon improvement of the conditions [7, 14].

### Quiescence Signals

Several pathways are responsible for the maintenance of quiescence in cancer cells. For example, the TGF-beta pathway has been reported to induce and maintain quiescence by inhibiting the cell cycle. The Notch signalling pathway also possesses the ability to impose a state of non-proliferation, in this case, within the tumour microenvironment. These are extremely crucial pathways in inducing long-lasting dormant cells [15, 19].

### Stress Response Pathways

Cancer cells will activate stress-related response in order to survive when the conditions are hostile. For instance, the p38 MAPK pathway becomes activated under environmental stress and can induce dormancy through the inhibition of cell growth and proliferation [8, 22].

### Tumour Heterogeneity

Not all cells in a tumour will enter or exit dormancy at the same time, and the tendency to enter dormancy can also vary considerably between them. This type of heterogeneity renders dormant cells challenging to treat because different subpopulations will respond in varying ways to environmental cues and therapeutic interventions [16, 3].

On the other hand, reactivation of dormant cancer cells can be influenced by a variety of factors, ranging from changes in the local microenvironment to genetic or epigenetic alterations:

### Microenvironmental Activation Triggers

Dormancy can be disrupted due to tumour microenvironment by alterations. For example, increased angiogenesis or even modifications in the immune system can provide quiescent cells with the stimulus necessary for re-initiation of cell growth and proliferation [1, 10, 20].

### Genetic or Epigenetic Activation Triggers

Dormancy is terminated by genetic mutations or epigenetic changes that rebalance proliferation and quiescence. These changes have a primary function in determining the duration of dormancy and the timing of reactivation [11, 13].

## 3. Experimental setting

The experimental setting that our model replicates was developed in [4]. Namely, the authors perform a series of measurements designed to investigate the spatiotemporal progression of metastasis in the lung, particularly focusing on the survival, dormancy, and (in-)efficiency of metastatic cancer cells. Here’s a detailed description of the experiment:

The researchers conducted an experiment using a mouse model to study the temporal and spatial progression of lung metastasis. They began by intravenously injecting B16F10 murine melanoma cells, labelled with a fluorescent dye, into mice. This allowed for the cells to circulate through the bloodstream and naturally arrest in the lungs, mimicking the early stages of metastasis.

Following the injection, the mice were sacrificed at various time points, and their lungs were examined to track the progression of the cancer cells. Through fluorescence microscopy, the researchers could observe whether the cells remained dormant, transitioned to active proliferation, or formed visible metastases of varying sizes. The use of microsphere markers allowed for precise quantification of cell survival, dormancy, and the growth of metastases over time.

This information was compiled in a flow chart, repeated here in Table 1 is fundamental in understanding the sequence of events that determine the fate of these cancer cells. It outlines the stages of initial arrest, extravasation, the decision between dormancy and proliferation, and the transitions in metastasis size—from single cells to small clusters, and eventually to macroscopic tumours.

**Table 1.** The metastatic progression of the cancer cells in the first 14 days after injection expressed relative to the number of initially injected cancer cells. Shown also the percentage of cancer cell loss; data from [4].

| Time<br>[d] | Solitary<br>cells<br>[%] | Metastases |  |  | Total<br>loss<br>[%] |
| --- | --- | --- | --- | --- | --- |
| | | Small<br>< 80 $\mu\text{m}$<br>[%] | Medium<br>80 – 300 $\mu\text{m}$<br>[%] | Large<br>> 300 $\mu\text{m}$<br>[%] | |
| 1/24 | 98 | 0 | 0 | 0 | 2 |
| 1 | 83 | 0 | 0 | 0 | 17 |
| 3 | 74 | 0 | 0 | 0 | 26 |
| 4 | 24.5 | 8.2 | 0.1 | 0 | 67.2 |
| 6-8 | 15 | 10.7 | 3.1 | 0.4 | 70.8 |
| 12-14 | 3.5 | 1.8 | 5.1 | 5.8 | 83.8 |

The chart reveals that dormancy is a critical bottleneck in the metastatic process, where many cancer cells, despite successfully arresting in the lung and extravasating into tissue, fail to transition from dormancy to active growth. This stage is pivotal as it dictates whether cells remain as dormant single entities or proceed to form larger, detectable metastases. Moreover, even when cells escape dormancy, their ability to proliferate and increase in size varies greatly, with many forming only small clusters that do not reach clinically significant sizes. These insights underscore the importance of targeting both dormant cells and the transitions to larger metastasis sizes in therapeutic interventions, as these are crucial points where the metastatic process could be effectively disrupted.

## 4. Mathematical Modelling

The mathematical model that we develop here, provides a framework to qualitatively and quantitatively describe the complex interactions and dynamics governing the survival, dormancy, and proliferation of metastatic cells, allowing for a deeper understanding of the above experimental observations.

This model that we propose describes the process of cancer cell arrest in the lung tissue, focusing on cell death in the blood stream, extravasation in the lung tissue, cancer cells dormancy and re-activation, and cancer cell proliferation and death.

These processes are described in step-by-step process:

### 4.1 In the bloodstream

Upon entering the circulatory system, the initial population of *N* ^init^ = 2× 10^5^ B16F10 melanoma cells injected in the caudal vein (tail vein), is directed by the blood flow primarily towards the lung tissue. Due to stresses in the blood, loss of survival signals, and the activity of the immune system, a number off the Circulating Tumour Cells (CTCs) is destroyed while in the blood stream.

Observations in [4] indicate that no CTCs are detected in the arterial and venous vessels 1 h post-injection, suggesting that the survival and extravasation of cells from the circulatory network into the lung tissue are highly efficient. Furthermore, these observations indicate that 98% of the CTCs survive the blood stream and arrest in the microvasculature of the lungs.

These results mean that the number of cells surviving and arresting in the tissue *t* = 1 h post-injection is:

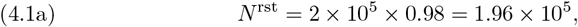

consequently the number of CTCs lost in circulation is:

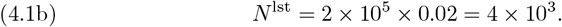

An alternative approach, that takes into account the migration time of CTCs through the bloodstream, could be obtained through a decay ODE of the form:

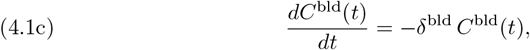

where *C*^bld^ the population size of of CTCs in the blood stream and *δ*^bld^ their— assumed constant—*death rate*. The solution to this equation is exponentially decaying:

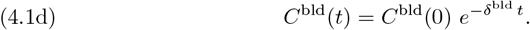

We take one more step and, for the sake of simplicity of the calculation of the rate *δ*^bld^, we assume that at time *t* = 1 h post-injection, a single cancer cell is left in the circulatory system; this implies, from (4.1d), that

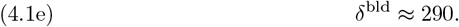

### 4.2 Dormancy

Post-extravasation, the disseminated tumor cells (DTCs) arrest in the lung occurs within a short period time. We assume that after arrest, each of the DTCs enters a dormancy phase that lasts for a period of time 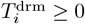, where *i* = 1 … *N* ^rst^. During this period, the cancer cells remain in a quiescent state, during which they do not proliferate.

To model the reactivation times, 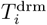, we consider a number of probability distributions that, at least initially, are theoretically justified. In a subsequent stage, Section 5, we fit these probability distribution to the experimental data of Table 1, identify their corresponding parameters, and perform comparative analysis.

### Suitable distribution functions

The current section provides an overview of the chosen probability distributions, including the formulas for their corresponding probability density functions (pdfs).

#### Normal distribution

This distribution is useful when reactivation times are expected to cluster around a central value, with equal likelihood of deviation on either side of the mean.

For *µ* and *σ* the mean and standard deviation:

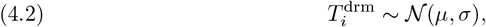

with the corresponding probability distribution function (pdf)

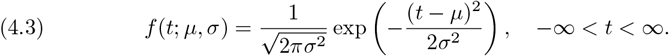

#### Logistic distribution

This distribution is suitable for modelling reactivation times with broader variability, capturing early and late reactivation. It is similar to the normal distribution but with heavier tails.

For *U* ∼ 𝒰 (0, 1) a uniform random variable, and *σ* the spread, the reactivation time reads:

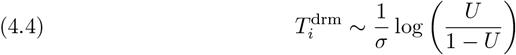

with pdf

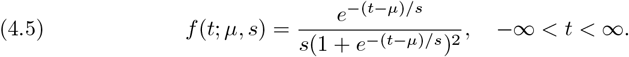

#### Gamma distribution

This distribution is useful for modelling reactivation that depends on a series of events or stages. For *k* and *θ* the shape and scale parameters:

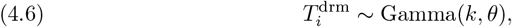

whose pdf is

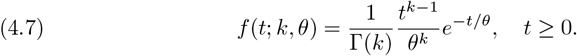

#### Weibull distribution

This distribution is ideal for scenarios where the likelihood of reactivation changes over time, such as increasing risk with time.

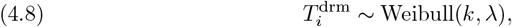

For *k* and *λ* the shape and scale parameters:

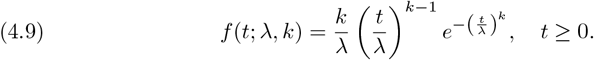

with the corresponding pdf

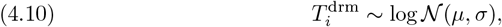

#### Log-Normal distribution

This distribution is appropriate for scenarios where early reactivation is rare, but the likelihood increases over time.

For *µ* and *σ* the mean and standard deviation:

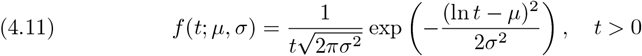

and the pdf is given by

#### Pareto distribution

This distribution is best for modelling scenarios where a majority of cells reactivate quickly, but a few may remain dormant for much longer.

If *U* ∼ 𝒰 (0, 1), (uniform distribution) and *α* and *t*_*m*_ the shape and scale parameters, then:

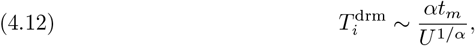

with the corresponding pdf

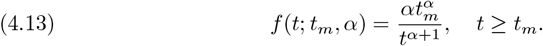

#### Gompertz distribution

This distribution is useful when reactivation becomes more likely as time progresses, reflecting a time-dependent increase in risk.

For *U* ∼ *u* (0, 1), and *b* and *η* the shape and location parameters:

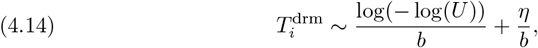

with the corresponding pdf

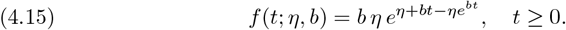

#### Inverse Gaussian distribution

This distribution is ideal for scenarios where reactivation time is influenced by random molecular events, providing a right-skewed distribution.

For *µ* and *λ* the mean and shape parameters:

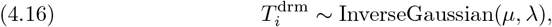

with the corresponding pdf

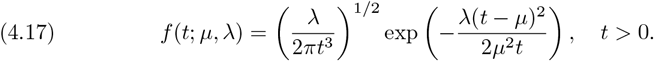

### Unsuitable distribution functions

Certain distributions are not well-suited to represent re-activation time. The following section will explore why these distributions fall short in this context.

#### Exponential distribution

The exponential distribution is often used to model time-to-event processes with a constant *hazard rate*, where the likelihood of an event occurring is the same at every time point. However, dormancy and reactivation times often involve changing probabilities over time, influenced by various biological factors. As such, the exponential distribution is not suitable for capturing these dynamic, time-dependent processes.

#### Poisson distribution

The Poisson distribution is typically used to model the number of discrete events occurring within a fixed interval of time. However, dormancy and reactivation times are continuous variables that describe the duration until a specific event, such as reactivation, occurs. As such, the Poisson distribution is not appropriate for modelling these continuous time-to-event processes.

#### Beta distribution

The Beta distribution is primarily used to model variables that are constrained within a fixed interval, typically [0, 1], such as probabilities or proportions. Dormancy and reactivation times, however, are continuous variables that can extend over an indefinite range, making the Beta distribution ill-suited for accurately representing these time-to-event processes.

Further distributions with similar characteristics, e.g. Uniform, Bionomial, Geometric, Bernoulli distributions, are equally not suited for this model, cf. [6].

### 4.3 Death and Proliferation in the lung

After reactivation, the previously dormant DTCs begin to proliferate in the lung tissue. We model this process as a continuous growth mechanism, without considering the cells’ relative position within the tissue or their proximity to blood vessels, and assuming a constant proliferation rate *r*^prl^ once reactivated.

Consequently, each reactivated DTC *i* = 1 … *N* ^rst^ gives rise to a (micro-)tumour of size *C*_*i*_(*t*), which grows according to the following ODE:

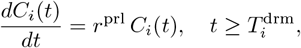

where 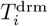 is the time at which the *i*-th cell reactivates.

Additionally, we assume that all dormant and active cancer cells experience death, modelled with a constant death rate *δ*^tss^. The combined model that accounts for both proliferation and death is represented by the following ODE:

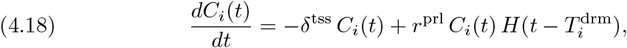

where 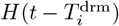 is the *Heaviside step function*, defined as:

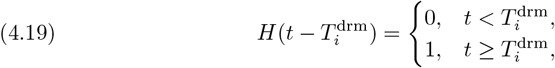

ensuring that proliferation only starts at 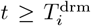, where 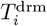 is the reactivation time of the *i*-th cancer cell.

### 4.4. Micro-tumour sizes

Upon reactivation, every DTC, indexed by *i* = 1 … *N* ^rst^, begins to proliferate, leading to the formation of a cluster of descendant cancer cells. In our model, the growth of each individual cluster is governed by the proliferation-death dynamics described in equation (4.18).

The size of these cell clusters is an important feature in assessing the progression of the metastatic tumour growth. Hence, we represent by 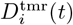 the diameter of the cluster initiated by the *i*-th DTC at time *t* after it has exited dormancy. Based on their diameters, we categorize the clusters into three distinct groups:

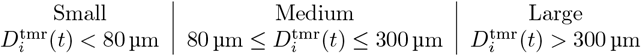

These size categories are selected as they reflect key biological stages in cluster development:

- Small clusters represent early growth stages and are typically undetectable through conventional imaging techniques.
- Medium clusters are likely detectable and indicate that proliferation has progressed significantly.
- Large clusters suggest the potential for further malignant progression, with possible implications for metastasis and clinical detection.

Moreover, through the growth dynamics modelled by (4.18), we can simulate the transition of cell clusters across these categories over time; this contributes to a better understanding of the metastatic spread.

## 5. Model parametrisation

The death rate *δ*^bld^ of the CTCs in the blood-stream, used in (4.1d), can be estimated using the measurement data of the successfully arrested cells in the lung *N* ^rst^ (4.1a), reported in [4] and the population size growth estimate (4.1d), to read

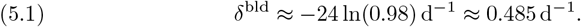

The doubling time for B16F10 cells has been reported with some variation in the literature. Namely, [18, 23, 21] reported doubling times of 20.1 h, 14.2 h, and 24 h, respectively, whereas [5] found a doubling time of 17.2 h. These differences are, according to [5], likely due to variations in the composition of the growth media used in each study. For the needs of our modelling, we convert these doubling times *T* ^dbl^ to proliferation rates *r* through the formula:

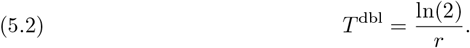

Accordingly, the calculated growth rates are *r*_1_ ≈ 0.0403 h^*−*1^, *r*_2_ ≈ 0.0345 h^*−*1^, *r*_3_ ≈ 0.0488 h^*−*1^, and *r*_4_ ≈ 0.0289 h^*−*1^. There is no particular reason to choose one of these rates over the others, hence we compute and compare their corresponding arithmetic and geometric mean values:

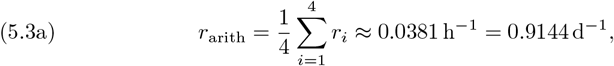

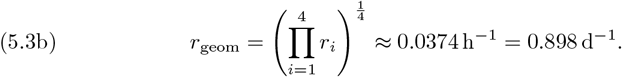

Since the growth rates are multiplicative in nature, we opt for the geometric mean as a more accurate growth rate; hence the proliferation rate *r*^prol^ that we employ in our growth model (4.18) is

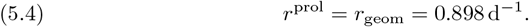

### 5.1. Comparison with experimental data

For the parameters that cannot be inferred directly—or indirectly as above—we assess them through the available data using specialized parameter estimation techniques.

These rely on the relative error, denoted as *E*_rel_, which compares the predictions of the mathematical model predictions *M* with the experimental/measurement data *D*, as summarized in Table 1. More specifically, the relative *L*_1_-error is defined through

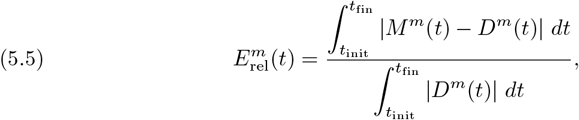

where *t*_init_ = 1*/*24h, *t*_fin_ = 14h and where the index *m* = 1 … 5 refers to the various metastases categories, cf. Table 1, and *t* the corresponding time of the measurements. We approximate these integrals using the trapezoidal rule over the discrete time points *t*_*i*_ ∈ {1*/*24, 1, 3, 4, 7, 14}h.

Once the relative error 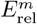 has been computed for all time points *t*_*i*_ and metastases categories, the overall/final value of the relative error is determined as the Root Mean Square (RMS) of the individual error values 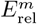:

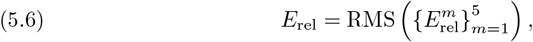

with the RMS formula is given by

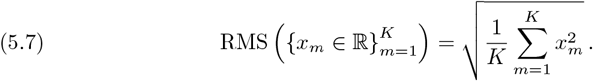

### 5.2 Parameter estimation

The parameter estimation technique that we employ is the Particle Swarm Optimization (PSO). PSO is a population-based optimization algorithm inspired by the collective behaviour of biological systems. It was first introduced in [12] and is primarily used to identify the global minimum/maximum of a function in multi-dimensional spaces. It is suited for continuous, nonlinear and potentially non-differentiable optimization problems.

PSO employs a large swarm of potential solutions of the optimisation problem. This swarm searches for the optimal solution by moving through the parameter by combining the information attained by every swarm particle individually and by the swarm as a whole. While moving through the parameter space, every particle of the swarm is aware of four pieces of information: its current location in the parameter space, its current velocity, the optimal position it has found during its exploration, and the best solution found by any particle in the swarm.

After initialisation with random position and velocity in the parameter space, the movement of every particle *i* is dictated by the formula

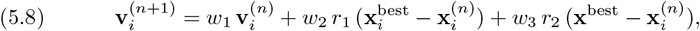

where *n* ∈ ℕ represents the iteration step of the method and where 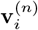 it’s current velocity, 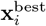 its personal optimal position, x^best^ the global optimal position found by any particle of the swarm. The weights *w*_1_, *w*_2_, *w*_3_ control the momentum, the memory and the influence of the global best position respectively. Furthermore, *r*_1_ and *r*_2_ denote uniformly distributed random values in [0, 1] responsible for the introduction of randomness/stochasticity in the movement of the particles.

Once the “new” velocity 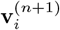 of the particle has been calculated, it’s position is adjusted according to:

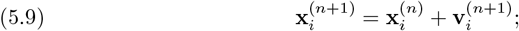

accordingly, the error function is calculated in this new position and the formula (5.8) is employed yet again and the process is repeated.

All implementations, simulations, and visualisations were conducted in [17].

## 6. Results

The PSO parameter estimation method described in Section 5.2 was applied for the death rate of cancer cells in the tissue and the two parameters of the probability distribution over the bounded domain Ω = [0, 10] × [0, 48] × [0, 24], respectively. Due to the lack of prior biological information for the bounds of these unknown parameters, the choice of domain Ω was based on extensive numerical experimentation, which indicated that expanding the domain further would not yield additional insights.

The results of the PSO are shown in Table 2, where the RMS relative *L*_1_ error (5.5) is presented along with the estimated death rate and the two corresponding parameters of the probability distributions.

**Table 2.** Results of the PSO parameter estimation method Section 5.2 applied to the various probability distributions considered in Section 4.2. The table includes the calculated error between model predictions and experimental data in Table 1, the estimated death rate for cancer cells in tissue, and the two controlling parameters for each distribution, also estimated. Notably, the death rate remains nearly invariant across different distribution choices.

| distribution | error | death rate | param. 1 | param. 2 |
| --- | --- | --- | --- | --- |
| Normal | 0.4101 | 0.2095 | 11.1644 | 4.1228 |
| Logistic | 0.4452 | 0.2362 | 10.9061 | 2.8597 |
| Gamma | 0.3720 | 0.2122 | 3.7747 | 3.2615 |
| Weibull | 0.3816 | 0.1961 | 14.2732 | 2.3760 |
| Log-Normal | 0.3628 | 0.2057 | 2.4833 | 0.6566 |
| Pareto | 0.2917 | 0.2122 | 4.6316 | 0.9665 |
| Gompertz | 0.4371 | 0.2406 | 4.6326 | 0.3966 |
| Inv. Gaussian | 0.3608 | 0.2087 | 17.2808 | 23.7758 |

The first remark is related to the death rate parameters, which show a remarkable consistency across all distributions i.e. 0.2095, 0.2362, 0.1950, 0.1961, 0.2576, 0.2122, 0.2406, 0.2019 with an average 0.2186. This consistency across different dormancy distributions strongly suggests that the estimated average death rate 0.2186 may closely approximate the true biological rate.

In Figure 1, we present the model predictions of the various tumour sizes against the experimental data, for each of the distributions and for the appropriate parameters. We order the graphs in an ascending error order and it even becomes visually clear that the fitness deteriorates significantly with the error. To gain, hence, a better insight into the distinct characteristics and suitability of the fit of the various distributions, we have evaluated several statistical measures of these distributions; their mean, variance, skewness, kurtosis, and cross-referenced them against the previously calculated relative error. These results are discussed below and are summarized in Table 3.

**Table 3.**
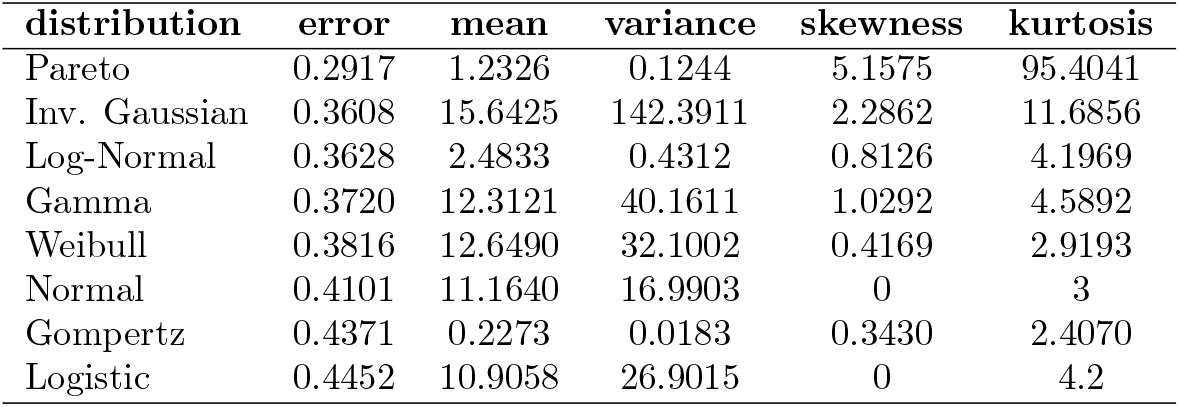
Statistical metrics for the probability distributions, including the calculated error between model predictions and experimental data. The table includes the mean, variance, skewness, and kurtosis for each one of the distributions. The suggestion is that those distributions with higher skewness and kurtosis may offer a more accurate fit for modelling re-activation times.

**Fig. 1.**
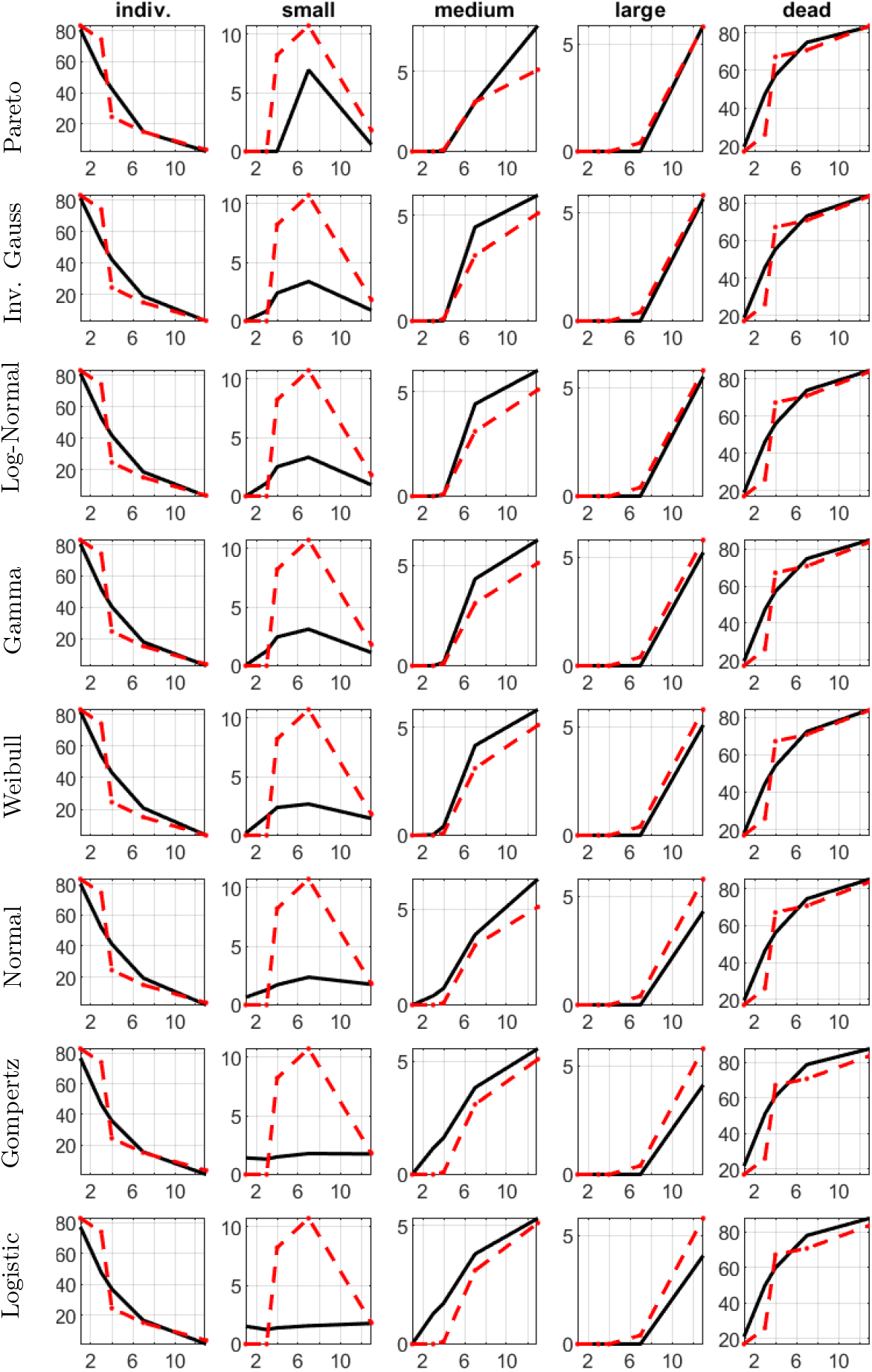
Comparison of model predictions (black solid line) and experimental data (red dashed line), for each one of the tested distributions. Each row represents a different distribution and each column a different tumour size: individual cells, small clusters (< 80 µm), medium clusters (80 ™ 300 µm), large clusters (> 300 µm), and overall dead tumour cells. The distributions are ordered row-wise according to increasing fitness error, cf. Table 2; it is evident that, from top-to-bottom, the fit between the model predictions and the experimental data, deteriorates.

Among the distributions considered, Pareto and Inverse Gaussian exhibit the lowest errors, and are coupled with high skewness and kurtosis. The high skewness and kurtosis values suggest a high degree of asymmetry and long tails, making them particularly suitable for capturing rare events or extreme values (outliers) in the data. Gamma and Weibull, exhibit slightly higher errors, moderate skewness, and lower kurtosis. While they still have positive skewness and higher kurtosis than the normal distribution, These two distributions result in a somewhat skewed but less heavy-tailed structure compared to Pareto and Inverse Gaussian. This makes them suitable for data that is moderately right-skewed but with fewer extreme values. Log-Normal, shows moderate positive skewness and higher kurtosis compared to the normal distribution. It performs comparably to the Gamma and Inverse Gaussian distributions in terms of error, though not as effectively as the Pareto distribution. In contrast, Normal and Logistic distributions display low skewness and kurtosis values common for symmetric distributions. Their higher errors suggest limitations in fitting highly skewed data with extreme values, as they are less capable of modelling asymmetry and heavy tails.

These results strongly indicate that the experimental data 1 represent cancer cell reactivation times with a right-skewed distribution and a pronounced tail extending toward longer durations. This implies that some reactivation times may be extreme events, and, hence, selecting models that accommodate this asymmetry and longtailed behaviour provide a more faithful representation of the reactivation times.

These remarks are reflected in Figure 1, where the model predictions for each of these distributions are plotted against the experimental data.

## 7. Conclusion

Understanding the processes of cancer cell reactivation and cancer cell duration dormancy are very important in cancer research, as dormancy increases the likelihood of metastasis and influences treatment outcomes. Identifying the distribution of dormancy durations provides valuable insights into the frequency of reactivation events and the survival times of dormant cancer cells under various conditions.

In this paper, we have examined the performance of cancer growth models—comprised of cancer cell death death, dormancy reactivation, and proliferation of cancer cell in the tissue—with a particular focus on the study of various statistical distributions in modelling cancer cell reactivation times. These were cross-referenced against experimental data in order to identify models that could effectively capture the asymmetry and variability observed in dormancy durations and the subsequent growth of cancer metastases. This evaluation aimed to identify not only the best-fit distribution but also those that are biologically interpretable.

During this process, we observed that the estimated death rate was consistent across all distributions, indicating that this parameter may be robust to the choice of a dormancy model. This stability further is a further indication of the biological relevance of the estimated death rate, providing confidence that it approximates the actual survival dynamics of cancer cells in dormancy.

The most effective distributions were the Pareto and Inverse Gaussian distributions, exhibiting the lowest fitness errors among all tested distributions (cf. Table 2). These distributions exhibit heavy-tailed behaviour and asymmetric skewness and are biologically relevant for rare and prolonged dormancy periods in cancer cells. On the other hand, distributions such as Gompertz and Logistic were less effective in capturing the dormancy periods and tumour sizes, because, as we argue, their mathematical structure did not align well with the biological properties of dormancy (cf. Figure 1).

Future extensions of this work should aim at expanding the framework that we have proposed here by introducing multiscale models that consider cell-cell and cell-tissue interactions. Furthermore, more appropriate and extensive data, extracted under different scenarios and animals are crucial. This would not help validate the proposed statistical distributions and also enhance the predictive capabilities and the biological applicability of the models.

## References

[1] J. A. Aguirre-Ghiso. Models, mechanisms and clinical evidence for cancer dormancy. Nature Reviews Cancer, 7(11):834–846, 2007.

[2] D. Barkan, D. El-Ashry, and A. F. Chambers. Metastatic growth from dormant cells induced by a col-i-enriched fibrotic environment. Cancer Research, 70(14):5706–5716, 2010.

[3] Philippe L Bedard, Aaron R Hansen, Mark J Ratain, et al. Tumour heterogeneity in the clinic. Nature, 501(7467):355–364, 2013.

[4] M. Cameron, E.E. Schmidt, N.A. Kerkvliet, K.V. Nadkarni, V.L. Morris, A.C. Groom, A.F. Chambers, and I.C. Macdonald. Temporal progression of metastasis in lung: cell survival, dormancy, and location dependence of metastatic inefficiency. Cancer research, 60 9:2541–6, 2000.

[5] C. Danciu, A. Falamas, C. Dehelean, C. Soica, H. Radeke, L. Barbu-Tudoran, F. Bojin, S. C. Pinzaru, and M. F. Munteanu. A characterization of four b16 murine melanoma cell sublines molecular fingerprint and proliferation behavior. Cancer Cell International, 13:75, 2013.

[6] M. H. DeGroot and M. J. Schervish. Probability and Statistics. Pearson, 4th edition, 2012.

[7] G. Flügen, A. Avivar-Valderas, Y. Wang, M. R. Padgen, J. K. Williams, A. R. Nobre, V. Calvo, J. F. Cheung, J. J. Bravo-Cordero, D. Entenberg, J. Castracane, V. Verkhusha, P. J. Keely, J. Condeelis, and J. A. Aguirre-Ghiso. Phenotypic heterogeneity of disseminated tumour cells is preset by primary tumour hypoxic microenvironments. Nature Cell Biology, 19(2):120–132, 2017.

[8] H. Gao, G. Chakraborty, and Adrian Lee-Lim. Multi-organ site metastatic reactivation mediated by non-canonical wnt signaling in human breast cancer cells. Nature Cell Biology, 14(7):748–756, 2012.

[9] C. M. Ghajar. Metastasis prevention by targeting the dormant niche. Nature Reviews Cancer, 15(4):238–247, 2015.

[10] C. M. Ghajar, H. Peinado, and H. Mori. The perivascular niche regulates breast tumour dormancy. Nature Cell Biology, 15(7):807–817, 2013.

[11] S. A. Joosse, T. M. Gorges, and K. Pantel. Biology, detection, and clinical implications of circulating tumor cells. EMBO Molecular Medicine, 7(1):1–11, 2014.

[12] J. Kennedy and R. Eberhart. Particle swarm optimization. In Proceedings of ICNN’95-International Conference on Neural Networks, volume 4, pages 1942– 1948. IEEE, 1995.

[13] C. A. Klein. Parallel progression of primary tumours and metastases. Nature Reviews Cancer, 9(4):302–312, 2009.

[14] X. Lu and Y. Kang. Hypoxia and hypoxia-inducible factors: master regulators of metastasis. Clinical Cancer Research, 16(24):5928–5935, 2010.

[15] S. A. Mani, W. Guo, M. J. Liao, E. A. Eaton, A. Ayyanan, A. Y. Zhou, F. Brooks, M. Reinhard, C. C. Zhang, M. Shipitsin, L. L. Campbell, K. Polyak, C. Brisken, J. Yang, and R. A. Weinberg. The epithelial-mesenchymal transition generates cells with properties of stem cells. Cell, 133(4):704–715, 2008.

[16] An. Marusyk, V. Almendro, and Kornelia Polyak. Intra-tumour heterogeneity: a looking glass for cancer? Nature Reviews Cancer, 12(5):323–334, 2012.

[17] MATLAB. MATLAB version 9.13.0.2105380 (R2022b). The Mathworks, Inc., Natick, Massachusetts, 2022.

[18] T. Ohira, Y. Ohe, Y. Heike, E. R. Podack, K. J. Olsen, K. Nishio, M. Nishio, Y. Miyahara, Y. Funayama, and H. Ogasawara. In vitro and in vivo growth of b16f10 melanoma cells transfected with interleukin-4 cdna and gene therapy with the transfectant. Journal of Cancer Research and Clinical Oncology, 120(11):631– 635, 1994.

[19] T. Oskarsson, S. Acharyya, X. H. F. Zhang, S. Vanharanta, S. F. Tavazoie, P. G. Morris, R. J. Downey, K. Manova-Todorova, E. Brogi, and J. Massaguè. Breast cancer cells produce tenascin c as a metastatic niche component to colonize the lungs. Nature Medicine, 20(8):937–947, 2014.

[20] S.Y. Park and J. S. Nam. The force awakens: metastatic dormant cancer cells. Experimental & Molecular Medicine, 2020.

[21] M. A. Qarawi, S. Carrington, I. S. Blagbrough, S. H. Moss, and C. W. Pouton. Optimization of the mtt assay for b16 murine melanoma cells and its application in assessing growth inhibition by polyamines and novel polyamine conjugates. Pharmaceutical Pharmacology Communications, 3(5–6):235–239, 1997.

[22] M. S. Sosa, F. Parikh, and A. G.etalaand others Maia. Nr2f1 controls tumour cell dormancy via sox9-and rarβ-driven quiescence programmes. Nature Communications, 6:6170, 2014.

[23] A. Yerlikaya and N. Erin. Differential sensitivity of breast cancer and melanoma cells to proteasome inhibitor velcade. International Journal of Molecular Medicine, 22(6):817–823, 2008.

